# Complete genome screening of clinical MRSA isolates identifies lineage diversity and provides full resolution of transmission and outbreak events

**DOI:** 10.1101/522078

**Authors:** Mitchell J Sullivan, Deena R Altman, Kieran I Chacko, Brianne Ciferri, Elizabeth Webster, Theodore R. Pak, Gintaras Deikus, Martha Lewis-Sandari, Zenab Khan, Colleen Beckford, Angela Rendo, Flora Samaroo, Robert Sebra, Ramona Karam-Howlin, Tanis Dingle, Camille Hamula, Ali Bashir, Eric Schadt, Gopi Patel, Frances Wallach, Andrew Kasarskis, Kathleen Gibbs, Harm van Bakel

**Author notes:** **Correspondence to**: Harm van Bakel (H.v.B.), One Gustave L. Levy Place - Box 1498, New York, NY 10029-6574. These authors contributed equally to this work (Mitchell J Sullivan, Deena R Altman, Harm van Bakel). These authors are co-senior authors on this work (Kathleen Gibbs, Harm van Bakel).

## Abstract

Whole-genome sequencing (WGS) of *Staphylococcus aureus* is increasingly used as part of infection prevention practices, but most applications are focused on conserved core genomic regions due to limitations of short-read technologies. In this study we established a long-read technology-based WGS screening program of all first-episode MRSA blood infections at a major urban hospital. A survey of 132 MRSA genomes assembled from long reads revealed widespread gain/loss of accessory mobile genetic elements among established hospital- and community-associated lineages impacting >10% of each genome, and frequent megabase-scale inversions between endogenous prophages. We also characterized an outbreak of a CC5/ST105/USA100 clone among 3 adults and 18 infants in a neonatal intensive care unit (NICU) lasting 7 months. The pattern of changes among complete outbreak genomes provided full spatiotemporal resolution of its origins and progression, which was characterized by multiple sub-transmissions and likely precipitated by equipment sharing. Compared to other hospital strains, the outbreak strain carried distinct mutations and accessory genetic elements that impacted genes with roles in metabolism, resistance and persistence. This included a DNA-recognition domain recombination in the *hsdS* gene of a Type-I restriction-modification system that altered DNA methylation. RNA-Seq profiling showed that the (epi)genetic changes in the outbreak clone attenuated *agr* gene expression and upregulated genes involved in stress response and biofilm formation. Overall our findings demonstrate that long-read sequencing substantially improves our ability to characterize accessory genomic elements that impact MRSA virulence and persistence, and provides valuable information for infection control efforts.

## Introduction

Healthcare-associated infections (HAI) with methicillin-resistant *Staphylococcus aureus* (MRSA) are common, impair patient outcomes, and increase healthcare costs (Klevens et al. 2007; Grundmann et al. 2006). HAI MRSA is highly clonal and much of our understanding of its dissemination has relied on lower resolution molecular strain typing methods such as pulsed-field gel electrophoresis (PFGE), *S. aureus* protein A (*spa*) typing, and multilocus sequence typing (MLST) (DeLeo and Chambers 2009), and typically includes characterization of accessory genome elements that define certain lineages and are implicated in their virulence. Examples of the latter include the arginine catabolic mobile element (ACME), *S. aureus* pathogenicity island 5 (SaPI5) and the Panton-Valentine leukocidin (PVL)-carrying ϕSa2 prophage in the community-associated (CA) CC8/USA300 lineage (Diep et al. 2006; DeLeo et al. 2010). Molecular typing facilitates rapid screening but has limited resolution to identify transmissions in clonal lineages. Moreover, genetic changes can lead to alteration or loss of typing elements (Glaser et al. 2016; Montgomery et al. 2009; Uhlemann et al. 2014; Planet et al. 2015). As such, WGS has emerged as the gold standard for studying lineage evolution and nosocomial outbreaks (Köser et al. 2012; Price et al. 2013). Transmission analysis with WGS has been performed largely retrospectively to date (Azarian et al. 2015; Altman et al. 2014; Harris et al. 2010; Snitkin et al. 2012), although prospective screening with resulting interventions has also been described (Eyre et al. 2012; Köser et al. 2012).

In addition to lineage and outbreak analysis, WGS has furthered our understanding of *S. aureus* pathogenicity by delineating virulence and drug resistance determinants (Mwangi et al. 2007; Benson et al. 2014), including those related to adaptation to the hospital environment (Senn et al. 2016; Mwangi et al. 2007). Many of these elements are found in non-conserved ‘accessory’ genome elements that include endogenous prophages, mobile genetic element (MGE), and plasmids (Lindsay and Holden 2004; Sela et al. 2018). The repetitive nature of many of these elements means that they are often fragmented and/or incompletely represented in most WGS studies to date due to limitations of commonly used short-read sequencing technologies, curbing insights into their evolution (Sela et al. 2018). Recent advances in throughput of long-read sequencing technologies now enable routine assembly of complete genomes (Chin et al. 2013; Madoui et al. 2015) and analysis of core and accessory genome elements (Benson et al. 2014; Altman et al. 2014), including DNA methylation patterns (Fang et al. 2012), but these technologies have not yet been widely used for prospective MRSA surveillance.

Here we describe the results of a complete genome-based screening program of MRSA blood isolates. During a 16-month period we obtained finished-quality genomes for first blood isolates from all bacteremic patients. In addition to providing detailed contemporary insights into prevailing lineages and genome characteristics, we characterized widespread variation across accessory genome elements, impacting loci encoding virulence and resistance factors, including those commonly used as molecular strain typing markers. During an outbreak event in the neonatal intensive care unit (NICU) we performed additional sequencing of surveillance and clinical isolates, and were able to provide actionable information that discriminated outbreak-related transmissions, identified individual sub-transmission events, and traced the NICU outbreak origin to adult hospital wards. Finally, comparative genome and gene expression analyses of the outbreak clone to hospital background strains identified genetic and epigenetic changes, including acquisition of accessory genome elements, which may have contributed to the persistence of the outbreak clone.

## Results

### Complete genome surveillance reveals genetic diversity among clonal MRSA lineages

In order to characterize the genetic diversity of MRSA blood infections at The Mount Sinai Hospital (MSH) in New York City, US, we sequenced the first positive isolate from all 132 MSH inpatients diagnosed with MRSA bacteremia between fall 2014 and winter 2015. Single molecule real-time (SMRT) long-read length RS-II WGS was used to obtain finished-quality chromosomes for 122 of 132 isolates (92%), along with 145 unique plasmids across isolates (**Supplemental Table 1**). The remaining isolates were in one or more chromosomal contigs that could not be closed with available long-read sequencing data. We reconstructed a phylogeny from a multi-genome alignment (**Fig. 1A, S1A**), which identified two major clades corresponding to *S. aureus* clonal complexes 8 (CC8; 45·5% of isolates) and 5 (CC5; 50% of isolates) based on the prevailing multi-locus sequence types (ST) in each clade (ST5 and ST105/ST5, respectively). The CC8 isolates further partitioned among the endemic community-associated (CA) USA300 (80%) and the hospital-associated (HA) USA500 (20%) lineages (**Fig. 1B**), while CC5 isolates mainly consisted of USA100 (75·8%) and USA800 (15·2%) HA lineages (**Fig. 1C**). Overall, the phylogeny was consistent with major *S. aureus* lineages found in the NYC region and the US (Pardos de la Gandara et al. 2016).

**Figure 1.**
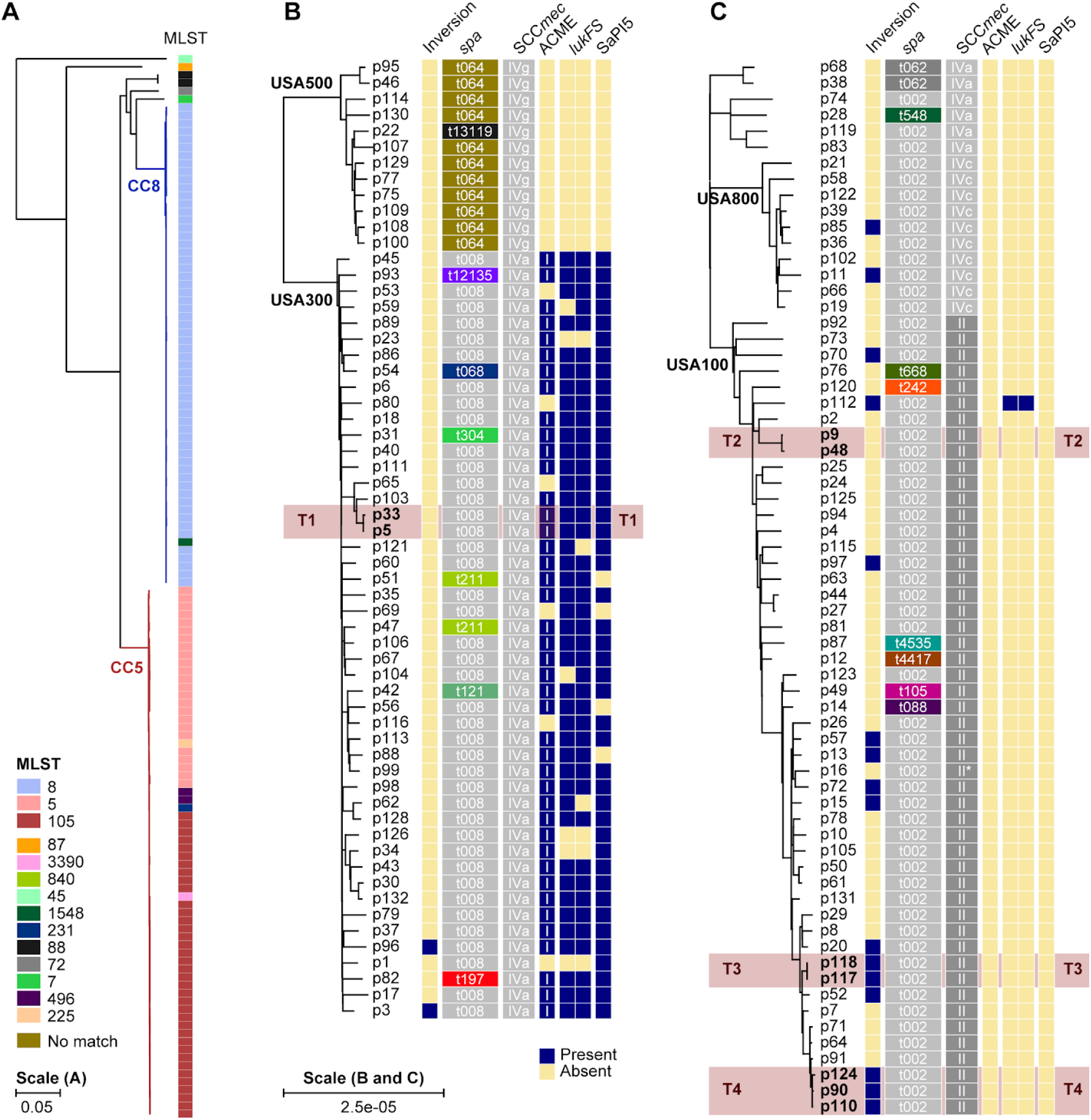
Phylogeny of MRSA bacteremia surveillance isolates. **A)** Maximum-likelihood phylogenetic tree based on SNV distances in core genome alignments of 132 primary MRSA bacteremia isolates. CC8 and CC5 clades are shaded in red and blue, respectively. Multilocus sequence types (MLST) for each branch are shown as coloured blocks, with a key at the bottom-left. **B)** Enlarged version of the CC8 clade from A. The isolate identifier is indicated next to each branch, together with blocks denoting the presence of large inversions (>250 kb), *spa* type, SCC*mec* type, and the presence (blue) or absence (yellow) of intact ACME, *lukFS* and SaPI5 loci. The ACME type is indicated in each box. The *lukFS* locus is represented by two blocks indicating the presence of *lukF* and *lukS*, respectively. **C)** Same as B, but for the CC5 clade. Asterisk indicates a spa type II isolate with an inserted element in the locus. Four transmission events between patients are highlighted in red and labeled T1 to T4. Scale bars indicate the number of substitutions per site in the phylogeny.

We next examined larger (>500 bp) structural variation that may be missed by short-read based WGS approaches (Copin et al. 2017; Altman et al. 2014; Harris et al. 2010). The multi-genome alignment indicated that between 80·8-88·9% of the sequence in each genome was contained in core syntenic blocks shared among all 132 genomes (**Supplemental Fig. 1A**). Another 9·5-16·8% was contained in accessory blocks found in at least two but not all genomes. Many of these accessory genome elements were lineage-specific and associated with prophage regions and plasmids (**Supplemental Fig. 1B**). Finally, 0·8-4·5% of sequence was not found in syntenic blocks and included unique elements gained by individual isolates. For example, a 32 kbp putative integrative conjugative element (ICE) carrying genes encoding proteins involved in heavy metal resistance (cadmium, cobalt, and arsenic) and formaldehyde detoxification was inserted after the *rlmH* gene in the USA800 isolate from p58 (**Supplemental Fig. 1C**). Similar arsenic resistance elements have been found in *S. aureus* isolated from poultry litter (Williams et al. 2006), which were linked to use of organic arsenic coccidiostats for growth promotion.

The extent of core and accessory genome variability impacted loci that are commonly used for molecular strain typing. Divergence from the dominant spa type was apparent in 8 (13-3%) of CC8 and 9 (13-6%) of CC5 lineage isolates. MLST loci were more stable in comparison with changes in 1-5% and 7-6% of isolates in each lineage, respectively. Notably, there were also widespread changes at ACME, PVL, and SaPI5 (**Fig. 1B**) in USA300 isolates, which are signature elements of this CA lineage (DeLeo et al. 2010; Diep et al. 2006) – 33-3% (16 of 48) either carried inactivating mutations or had partially or completely lost one or more elements (**Fig. 1B**). The multiple independent events of ACME, PVL and SaPI5 loss throughout the USA300 clade may reflect its ongoing adaptation to hospital environments, as these elements are typically absent in HA lineages. Interestingly, we found one case of a PVL-positive USA100 isolate (**Fig. 1C**) that may have resulted from homologous recombination between a ¦Sa2 and ¦Sa2 PVL prophage (**Supplemental Fig. 2**). Thus, complete genomes of MRSA blood isolates demonstrate the mobility of the accessory genome in ways that impact commonly used *S. aureus* lineage definitions.

The multi-genome alignment further identified large inversions spanning >250 kbp in 18 genomes (13-6%) (**Supplemental Fig. 1A**), which were much more common in CC5 (16 inversions) vs. CC8 lineages (2 inversions) (**Fig. 1**). The ends of these large inversion events were mainly (94%) located within distinct prophage elements that shared large (>10kb) regions of high sequence similarity (>99%), which meant that the exact cross-over points could not be identified. Notably, 11 inversions spanning ~1·15 Mb occurred between ΦSa1 and ΦSa5 in CC5 isolates and could only be resolved by using raw long-read data to phase the small number of variants that uniquely differentiated each prophage (**Supplemental Fig. 3**). Other inversions involved cross-overs between prophage pairs of ΦSa1, ΦSa3, ΦSa7, and ΦSa9. The chimeric prophages that resulted from the inversions consisted of new combinations of the two original prophage elements and contained all genes necessary to produce functional phages based on PHASTER (Arndt et al. 2016) analyses. Taken together, this suggests that prophage elements are common drivers of large inversion events in *S. aureus* that contribute to prophage diversity.

### Identification of transmission events among adults and an outbreak in the NICU

We next compared isolate genomes to identify transmissions between patients. We considered intra-host diversity and genetic drift in aggregate and set a conservative distance of ≤7 SNVs to define transmission events (see Methods). At this threshold we identified one USA300 and three USA100 transmissions involving six adults and three infants (**Fig. 1B-C**, labeled as T1-T4). Complete genome alignments for each event confirmed the absence of structural variants. In the USA300 transmission case (T1) the presumed index patient p5 was bacteremic with the same clone on two occasions ~3 months apart (**Supplemental Fig. 4**). The isolate obtained from the recipient (p33), who was later admitted to the same ward for 7 days at the time of collection. The USA100 isolates in transmission T2 were collected ~4 months apart and although the patients had overlapping stays, they did not share a ward or other clear epidemiological links (**Supplemental Fig. 4**). In transmission T3, both patients shared a ward for several days (**Supplemental Fig. 4**).

The final transmission involving 3 infants (T4) was part of a larger outbreak in the NICU, where positive clinical MRSA cultures from three infants within five weeks had prompted an investigation and consultation with the New York State Department of Health (NYSDOH). During four months an additional 41 clinical and surveillance cultures from 20 infants tested positive for MRSA, bringing the total to 46 isolates from 22 infants. Three further isolates were obtained from incubators and an IV box, from a total of 123 environmental swabs (2-4 %). Positive nasal surveillance cultures were also obtained from 2 out of 130 (1-5%) healthcare workers (HCWs) who had provided direct care to newly MRSA-colonized infants. The NYSDOH performed PFGE on 22 isolates, of which 14 patient and 3 environmental isolates had nearly indistinguishable band patterns (data not shown). This included p90 and p110 in transmission T4 (**Fig. 1C**; p125 was not tested). The USA100 (ST105) outbreak clone was resistant to fluoroquinolones, clindamycin, gentamicin and mupirocin, and susceptible to vancomycin, trimethoprim-sulfamethoxazole and doxycycline (**Supplemental Fig. 5, Supplemental Table 1**). This pattern was uncommon (18·2%) among USA100 isolates in our study and was therefore used as an initial screening criteria for cases. None of the HCW isolates matched the MLST or antibiogram of the outbreak clone and both staff members were successfully decolonized with nasal mupirocin and chlorhexidine gluconate (CHG) baths.

### Complete genome surveillance resolves outbreak origin and progression

During the outbreak we expanded our genomic screening program to include the first isolate of suspected outbreak cases. From day 354 onwards we obtained 23 additional complete genomes (**Supplemental Table 1**). Of these, 19 genomes from 16 infants and three environmental isolates matched the ST105 outbreak strain type, bringing the total to 22 outbreak genomes from 16 infants and the environment. We reconstructed a phylogenetic tree based on core genome alignments of all ST105 isolates in our study, which grouped all 22 isolates with matching antibiograms and/or PFGE patterns in one well-defined clade (**Fig. 2A**).

Surprisingly, this clade also contained 3 MRSA isolates obtained from adult bacteremia patients in other hospital wards prior to the first NICU case. The outbreak clade genomes were ≤15 SNVs apart, and the clade as a whole differed from other ST105 isolates by ≥41 SNVs. We therefore considered the 3 adult isolates to be part of a larger clonal outbreak that spanned 7 months. Based on the pattern of variants between outbreak genomes we could distinguish 4 distinct subgroups (**Fig. 2A, 2B**).

**Figure 2.**
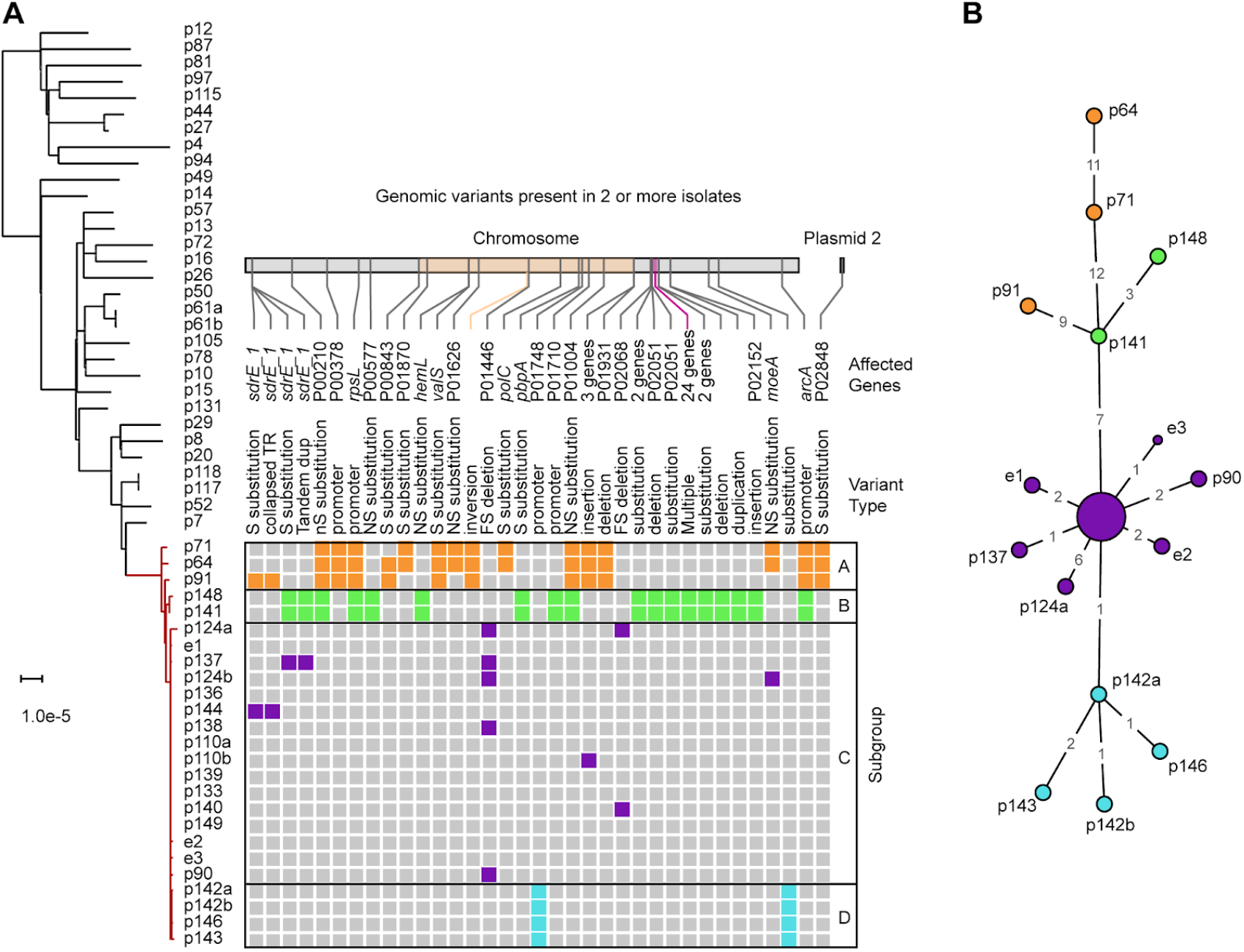
NICU outbreak subgroups and association with adult bacteremia patients. **A)** Maximum-likelihood phylogenetic tree based on SNV distances in core genome alignments of 31 ST105 primary bacteremia isolates (black) and 25 outbreak isolates (red). The scale bar indicates the number of substitutions per site. The patient (p) or environmental (e) isolate identifier is shown next to each branch (a/b suffixes indicate multiple isolates from the same patient). Variants present in two or more NICU outbreak isolates, derived from full-length pairwise alignments to the p133 genome, are shown as coloured boxes. Variants are colored according to outbreak subgroups inferred from common variant patterns, as indicated on the right. For each variant the genomic location, affected genes, and type of mutation is shown above the matrix. A 2 Mbp inversion in the adult isolates and a 2,411 bp region containing two substitutions and a deletion in subgroup B is highlighted in the location bar in orange and purple, respectively. **B)** Minimum spanning tree of the 25 outbreak isolates based on SNVs identified in their complete genome alignments. The 15 labeled nodes represent individual isolates. The larger central node corresponds to ten isolates with identical core genomes, which includes the p133 reference. Nodes are colored according to the outbreak subgroups shown in panel A. Numbers at edges represent core genome SNV distances.

We then used the available epidemiology and genomic data to reconstruct an outbreak timeline (**Fig. 3A**). The three initial adult cases had overlapping stays and shared wards, and their isolates clustered together in subgroup A. Several of the earliest clinical isolates from infants p141, p150, and p151 that coincided with the spread of the outbreak to the NICU were not available for genomic analysis (marked *X* in **Fig. 3A**). The missing isolate from p141 was susceptible to gentamicin and differed from the PFGE pattern of the outbreak clone by five bands. The other two missing isolates from p150 and 151 matched the outbreak clone antibiogram and were therefore considered to be part of the outbreak. Subsequent cases were identified by positive surveillance cultures on days 357-386 and their isolates clustered in subgroup C. All but one of the infants in this subgroup stayed in NICU room 2 before or at the time of culture positivity. The three positive environmental isolates were also obtained from this room, suggesting that a local bioburden lead to a high volume of colonized infants in a short time. Construction in the NICU and a resulting disruption of infection prevention practices was believed to play a role in the initial transmissions of MRSA.

**Figure 3.**
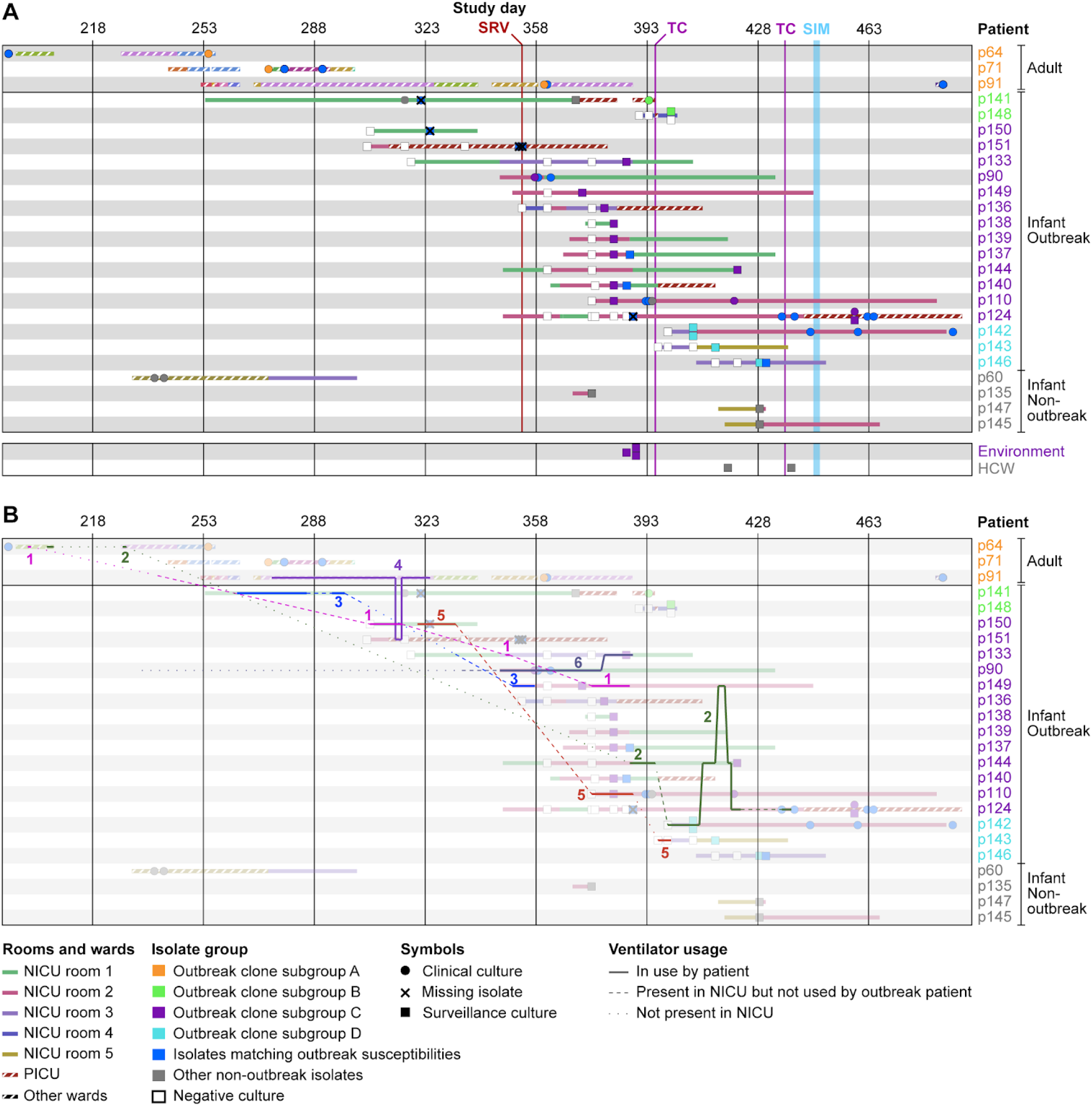
Timeline of the NICU outbreak. **A)** Overview of outbreak patient stays and isolates collected during the NICU outbreak. Rows correspond to patients with admission periods shown as horizontal bars. Solid fill patterns denote NICU stays and striped patterns indicate stays in other MSH wards. Fill colors correspond to NICU rooms (solid) or hospital wards (striped). Clinical or surveillance isolates collected during each stay are indicated by symbols, with a key shown below. Patient identifiers and isolate symbols are colored by outbreak subgroup. Timeline scale and key interventions are shown at the top. SRV - start of biweekly surveillance cultures; TC - terminal cleaning; SIM - *in situ* simulation. **B)** Same as A, but with ventilator movements between patients and locations overlaid as lines. Ventilators are numbered and shown in distinct colors. Solid lines correspond to periods that a ventilator was in use by an outbreak patient. Dashed lines indicate when a ventilator was present in the NICU but not used by an outbreak patient. Dotted lines indicate when a ventilator was not in use by an outbreak patient and not present in the NICU. Background colors are muted to facilitate tracking of ventilator movements.

The increase in new cases on surveillance prompted a terminal clean (TC) of the NICU on day 395. During this time, all infants were temporarily transferred to two different locations. Infant p148 who was colonized with the outbreak clone was placed across the hall from p141 in the pediatric intensive care unit (PICU). A positive surveillance culture in the same subgroup (B) as p141 was obtained for p148 shortly afterwards (**Fig. 3A**), suggesting that a transmission had occurred during the TC. New positive surveillance cultures were subsequently found for three additional infants (p142, p143 and p146). Each had been admitted after the TC and stayed in room 3 before or at the time of culture positivity. Their isolates comprised subgroup D, suggesting that the outbreak clone spread to this location from the closely related subgroup C linked to room 2 (**Fig. 2B, 3A**). Thus, each outbreak subgroup (A-D) was associated with a specific area (adult wards, PICU, and NICU rooms 2 and 3, respectively), indicating that location sharing was a dominant factor in the spread of the outbreak clone.

The continued transmissions after the first TC prompted *in situ* simulation and a second TC (**Fig. 3A**). The simulation efforts reinforced the importance of compliance to infection prevention strategies, patient cohorting, enhanced environmental disinfection, and limiting patient census to decrease bioburden (Gibbs et al. 2018). Only one new case (p124) was detected after the second TC. Infant p124 was located the PICU at the time of detection and based on the genomic profile (subgroup C) and earlier positive isolates, the transmission was believed to have occurred prior to the final TC and *in situ* simulation. As such, the workflow improvements were effective in halting the outbreak. The weekly surveillance cultures ended after three consecutive weeks of negative cultures on day 452. The last colonized patient was discharged two months later, and we did not detect the outbreak clone in our hospital-wide genomic screening program in the subsequent two years. While the majority of cases were positive by surveillance, there was morbidity related to the outbreak; five infants developed clinical infections, with three bacteremias, one pneumonia, and one surgical site infection. There were no deaths related to the outbreak.

### Role of ventilator sharing in the NICU outbreak

Location and HCW sharing could not account for the link between adult and pediatric cases, which were housed in different buildings and cared for by different HCWs. We focused on a potential role of ventilators in the outbreak based on the observations that: *i)* all NICU outbreak cases were on invasive or non-invasive ventilator support prior to culture positivity; *ii)* the three adult patients were ventilated for at least part of their hospitalizations; and *iii)* prior to identification of the NICU outbreak ventilators were shared between adult and pediatric wards. Ventilator exchange between units was discontinued after the first NICU cases were identified.

The ventilators present in the NICU during environmental surveillance tested negative for MRSA, but we could not rule out earlier contamination or contributions of other ventilators. Analysis of equipment usage logs and tracking data provided by the hospital’s real-time location system (RTLS) identified six units that were shared between outbreak cases (**Fig. 3B**, numbered 1-6). Ventilator 1 was briefly used by adult p64 and then transferred to several locations before it was moved into the NICU and later used by infant p150. The first NICU isolate that matched the outbreak clone by antibiogram was isolated from this patient soon after (**Fig. 3A, B**). Ventilator **4** was used by adult p91 several weeks before this patient developed bacteremia, except for a 2-day period when it was used by infant p151, shortly before the first NICU outbreak case (**Fig. 3B**). Infant p151 was in the neighboring PICU at this time and remained there until a positive surveillance isolate was obtained. Finally, ventilator 2 was used by adult p64 in two separate hospital visits, but was only moved to the NICU after the outbreak had already spread there.

Within the NICU, the sequential use of ventilator 6 by patient p90 and p133, the timing of their respective culture positivity, and the similarity of their isolate genomes, all supported a role for this ventilator in the transmission to p133. Likewise, ventilators 2 and/or 5 may have been a factor in the spread from room 2 (subgroup C) to room 3 (subgroup D), especially considering that both rooms were cleaned just prior to the transmission (**Fig. 3A**). Ventilator 5 may also have been a transmission vector from p150 to p110. Ventilator 3 was used by p141 and later by p149; however, it is unclear if it played a role in the outbreak, as the first two isolates obtained from p141 after ventilator 3 exposure did not match the outbreak. Altogether, the epidemiological and genomic data suggest that ventilators not only played a role in spreading the outbreak from adult wards to the NICU but were also a factor in subsequent sub-transmissions within the NICU.

### Mutations in the outbreak clone alter expression of virulence and persistence factors

Given the extended duration of the outbreak we next sought to identify genomic features that could have contributed to its persistence. A comparison of complete genomes found 42 non-synonymous or deleterious SNVs and indels in the outbreak clone that were not present in any of the ST105 hospital background strains, affecting 35 genes or their promoter regions (**Fig. 4A**). The products of these genes were primarily involved in nucleotide, amino acid and energy metabolism, as well as environmental signal processing and drug resistance. Several genes encoding cell wall proteins were also affected, including *gatD*, which is involved in amidation of peptidoglycan (Figueiredo et al. 2012).

**Figure 4.**
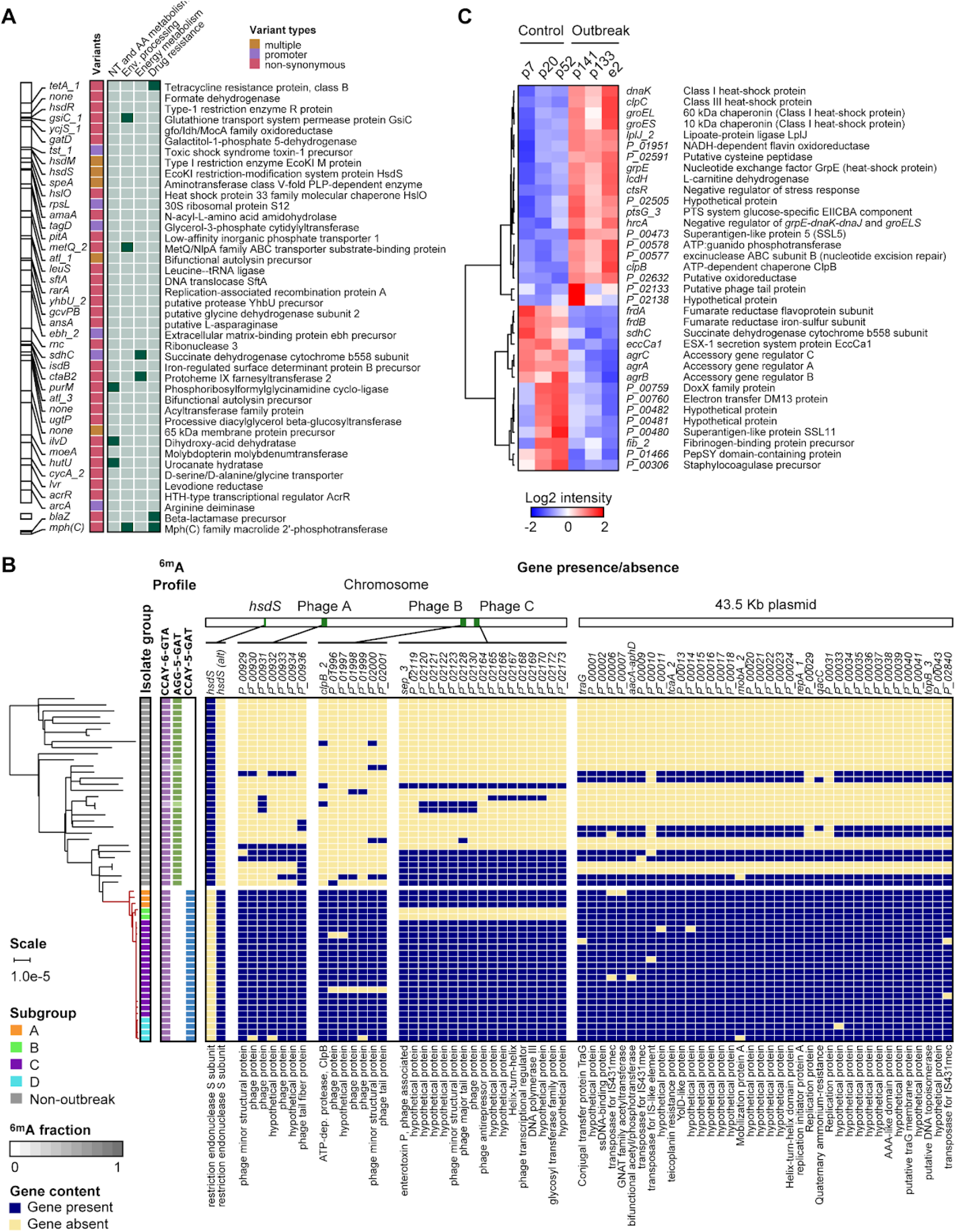
Differentiating features of the NICU outbreak clone compared to the USA100 background. **A)** Map of non-synonymous SNVs in genes and promoter regions that are unique to the outbreak clone. Gene identifiers or names are shown next to their genomic location. The SNV type is indicated by colors with a key shown at the top-right. KEGG pathways with two or more genes are indicated on the right (green boxes) and corresponding gene descriptions on the far-right. **B)** Pan-genome analysis of MLST105 isolates showing all genes present in the outbreak clone and absent from at least half of the non-outbreak isolates collected during our study. A maximum-likelihood phylogenetic tree based on SNV distances in core genome alignments is shown on the left with patient (p) or environmental (e) isolate identifiers. Changes in the ^6m^A methylation profile due to the *HsdS* recombination in the outbreak strain are highlighted in green/blue. Gene presence (yellow) or absence (red) is indicated in a matrix organized by genomic location (top). Gene names and descriptions are shown at the top and bottom of the matrix, respectively. See key on bottom left for more details. **C)** Hierarchical clustering of 35 genes with significant expression differences (FDR q<0·05) between three control and three outbreak strains. Columns correspond to control or outbreak isolates, with labels at the top. Gene names and descriptions are shown on the right. Color shades and intensity represent the difference in normalized log2 counts per million (CPM) relative to the average gene expression level, with a color key shown below.

Pan-genome analysis with Roary (Page et al. 2015) further revealed 71 genes exclusive to the outbreak strain or infrequently (<33%) present in other MLST105 isolates (**Fig. 4B**). Most of these genes were associated with three prophage regions and a 43·5 kbp plasmid. The additional genes in prophage A encoded only phage replication or hypothetical proteins. Among the genes in prophage B was an extra copy of *clpB*, which promotes stress tolerance, intracellular replication and biofilm formation (Frees et al. 2004). Prophage C included an extra copy of the sep gene encoding an enterotoxin P-like protein associated with an increased risk of MRSA bacteremia in colonized patients (Calderwood et al. 2014). The 43·5 kbp plasmid contained the mupirocin (*mupA*), and gentamicin (aacA-aphD) resistance genes (**Supplemental Fig. 5B**) that explained the distinct susceptibility profile of the outbreak clone. High-level mupirocin resistance (HLR) conferred by *mupA* has been linked to transmissions in previous studies (Hodgson et al. 1994; Udo et al. 2001). Pan-genome analysis also revealed a unique variant of the *HsdS* gene in the outbreak strain, which encodes the specificity subunit of a Type I restriction modification (RM) system. Closer examination revealed that a recombination event in one of the DNA recognition sites (**Supplemental Fig. 7**) changed a target recognition domain from the typical CC5 domains, “BD” (“AGG-5-GAT” present at 738 sites, overlapping 595 genes and 120 promoter regions) to “AD” (CCAY-5-GAT present at 304 sites, overlapping 287 genes and 15 promoter regions), resulting in altered genome-wide ^6m^A DNA methylation profiles compared to other ST105 isolates (**Fig. 4B**).

We reasoned that the (epi)genetic changes in the outbreak clone could alter gene expression patterns and provide further insights into the effects of these changes. We therefore compared the gene expression profiles of three representative outbreak isolates (i.e., cases) to the three most similar non-outbreak ST105 strains (i.e., controls) during late-log phase growth. The control strains shared the 43·5 kbp plasmid and most of the prophage elements with the outbreak strain and demonstrated similar growth characteristics (**Supplemental Fig. 7**). Differential gene expression analysis showed altered expression of 35 genes (**Fig. 4C**). Two of these genes were mutated in the outbreak clone; a SNP in promoter region of *sdhC* and a duplication of *clpB*. Methylation changes were found in six genes (17·1%), which was lower than the rate of 27·3% across all genes. Thus, most expression changes appear to be indirect results of (epi)genetic changes. Multiple upregulated genes in the outbreak clone encoded proteins involved in stress and heat shock responses. This included *clpB*, which was increased in copy number in the outbreak vs. control strains, but also *dnaK* and *clpC*, which have been linked to biofilm formation in *S. aureus* and adherence to eukaryotic cells (Singh et al. 2012; Chatterjee et al. 2005). Expression of the gene encoding staphylococcal superantigen-like protein 5 (SSL5) was also increased. SSL5 is known to inhibit leukocyte activation by chemokines and anaphylatoxins (Bestebroer et al. 2009). Among the downregulated genes, the *agrABC* genes of the accessory gene regulator (*agr*) locus stood out. *Agr* is the major virulence regulator in *S. aureus* (Novick 2003) and decreased *agr* function in clinical isolates is associated with attenuated virulence and increased biofilm and surface protein expression (Shopsin and Copin 2018). Taken together, the nature of the genetic and expression changes in the outbreak clone indicate they may have contributed to its persistence.

## Discussion

In this study we implemented a complete genome screening program at a large quaternary urban medical center, with the aim of tracking circulating clones, to identify transmission events, and to understand the genomic epidemiology of endemic strains impacting human health. To our knowledge, this is the largest set of clinical MRSA isolates from bacteremic patients to undergo complete genome assembly to date. The availability of complete genomes allowed us to precisely map all genetic changes between strains highlighting the presence of substantial structural variation in lineages that are commonly considered to be highly clonal. The extent of variation due to recombinations in prophages, mobilization of genetic elements, and large genomic inversions also impacted classical spa, MLST and signature virulence and resistance elements used in *S. aureus* molecular typing schemes. As such, the stability of these elements should be considered when using such schemes for lineage analysis. Complete reconstruction of outbreak genomes provided additional variation data to map sub-transmission events during a NICU MRSA outbreak. Finally, the combination of genetic and gene expression differences between the NICU outbreak clone and USA100 hospital background revealed genomic features that may have contributed to its persistence.

Much of the accessory genome variation occurred in prophage elements, further underscoring their importance in *S. aureus* genome organization (Xia and Wolz 2014). We also show that prophages are common drivers of large chromosomal inversions, with evidence of multiple independent events throughout the phylogeny. Inversions were much more frequent in CC5 and USA100, which may reflect higher similarity between endogenous prophages and/or the increased divergence between isolates in the USA100 lineage. Most inversions could only be resolved by long-read sequencing data, and our results, combined with our previous observations among MSSA isolates (Altman et al. 2018), suggest that prophage-mediated recombinations may be more frequent than previously appreciated. Indeed, one inversion event occurred in the outbreak clone during the spread from the adult wards to the NICU. The impact of genomic inversions on *S. aureus* and their clinical relevance is unclear and will require further study, but they likely explain the highly chimeric and mosaic structure of *S. aureus* prophages. Notably, non-reciprocal double break-and-join or long gene conversion events can facilitate sequence exchanges between prophages (Fortier and Sekulovic 2013). This could lead to the reshuffling of virulence genes and a wider horizontal spread as they become incorporated in phages with different host ranges.

Complete genome analysis of the outbreak clone revealed a pattern of genetic changes that matched patient locations, suggesting that transmission bottlenecks and local environmental contamination led to a unique genetic signature at each site. Some isolates and isolate subgroups were separated by >10 variants, which is relatively high considering a reported core genome mutation rate of 2·7-3·3 mutations per Mb per year (Harris et al. 2010; Young et al. 2012). This suggests that the outbreak may have originated from a genetically heterogeneous source, such as a patient with a history of persistent MRSA colonization that accumulated intra-host variants. It is also possible that the combination of selection pressures and transmission bottlenecks contributed to the diversification of the outbreak clone. Considering all available data, we think the most likely scenario is that the NICU outbreak originated from patient p64 and then spread to other adult patients through direct or indirect contact in shared wards. Ventilator 1, used by adult p64 and infant p150, is the most likely vector for entry into the NICU. Ventilator 4 may have provided a potential second entry route via p151, with subsequent transmissions to p141 and p148 (p151 and p141 had an overlapping stay in the PICU). Such a secondary introduction may explain why the p141 and p148 isolates were more distantly related to all other NICU isolates, but we were not able to test this scenario as the isolates from p151 were no longer available. All subsequent cases could be explained by location relative to other MRSA colonized patients or sharing of MRSA-exposed ventilators.

The outbreak strain genome differed from the hospital background by multiple mutations of core genes, as well as accessory gene gain and loss. Hundreds of genes were impacted by DNA methylation changes in the gene body or promoter regions, but such genes were depleted rather than enriched among differentially expressed genes. As such, the impact of the methylation changes on the outbreak clone (if any) was unclear. Nonetheless, a common theme among the genetic and expression changes was the relevance of genes involved in biofilm formation, persistence and quorum sensing. Although the collective impact of the mutations will require further investigation, we speculate that these changes may have contributed an increased persistence of the outbreak clone in the environment.

Complete genome data from our hospital-wide screening program provided key information for outbreak management that could not have been obtained by molecular typing. First, it provided conclusive differentiation of outbreak from non-outbreak isolates, which helped delineate the final case set and determine when the outbreak ended. Second, analysis of all genetic differences between outbreak cases allowed us to identify sub-transmissions and better understand the chain of events that led to each sub-transmission. Third, the availability of hospital-wide genomic surveillance data indicated that the NICU outbreak originated much earlier in unrelated adult wards in a different building and helped identify ventilators as likely transmission vectors.

There are some limitations to our study. Our genomic survey was limited to first positive single-patient bacteremias and transmission rates may be increased when considering non-blood isolates. Moreover, by sequencing single colony isolates we likely did not fully capture intra-host heterogeneity. Although such heterogeneity may be less common among bacteremias, we did encounter variation within some patients which was considered when establishing our transmission thresholds. Finally, while we believe that we have reconstructed the most likely transmission routes and vectors for the NICU outbreak, it is possible that other factors such as spread by HCWs and/or other vectors contributed as well.

In conclusion, we find that the application of complete genome sequencing in the clinical space provides significant benefits for infection prevention and control. In addition to providing contemporary data on the genomic characteristics of circulating lineages, directed intervention and containment of identified transmission events can help prevent further outbreak progression. Although our screening program was limited in scope to bacteremias, early detection of a transmission event between the adult and NICU ward could conceivably have allowed staff to intervene earlier. Completely finished genomes also provide the ability to identify unique elements of particular strains. Accumulating a larger repository of complete and unique genome references and variants associated with successful spreading strains may be key to future outbreak detection and prevention programs by providing high-resolution feature sets for prospective and retrospective data mining purposes.

## Methods

### Ethics statement

This study was reviewed and approved by the Institutional Review Board of the Icahn School of Medicine at Mount Sinai, and the MSH Pediatric Quality Improvement Committee.

### Case review

An investigation of the characteristics of the patients included review of existing medical records for relevant clinical data. Unique ventilator identification numbers and the real-time location system (RTLS) enabled mapping of ventilator locations over time.

### Bacterial isolate identification and susceptibility testing

Isolates were grown and identified as part of standard clinical testing procedures in the Mount Sinai Hospital Clinical Microbiology Laboratory (CML), and stored in tryptic soy broth (TSB) with 15% glycerol at −80°C. VITEK 2 (bioMérieux) automated broth microdilution antibiotic susceptibility profiles were obtained for each isolate according to Clinical and Laboratory Standards Institute (CLSI) 2015 guidelines and reported according to CLSI guidelines (Wayne 2015). Susceptibility to mupirocin was determined by E-test (bioMérieux) and susceptibility to chlorhexidine was tested with discs (Hardy) impregnated with 5 μl of a 20% chlorhexidine gluconate solution (Sigma-Aldrich). Species confirmation was performed with MALDI-TOF (Bruker Biotyper, Bruker Daltonics).

### DNA preparation and sequencing

For each isolate, single colonies were selected and grown separately on tryptic soy agar (TSA) plates with 5% sheep blood (blood agar) (ThermoFisher Scientific) under nonselective conditions. After growth overnight, cells underwent high molecular weight DNA extraction using the Qiagen DNeasy Blood & Tissue Kit (Qiagen, 69504) according to the manufacturer’s instructions, with modified lysis conditions. Bacterial cells were lysed by suspending cells in 3 μL of 100 mg/ml RNase A (Ambion, AM2286) and ten μL of 100 mg/ml lysozyme (Sigma, L1667-1G) for 30 minutes at 37°C, followed by incubation with Proteinase K for one hour at 56°C and two rounds of bead beating of one min each using 0·1mm silica beads (MP Bio) (Altman et al. 2014).

Quality control, DNA quantification, library preparation, and sequencing was performed as described previously (Altman et al. 2014). Briefly, DNA was gently sheared using Covaris G-tube spin columns into ~20,000 bp fragments, and end-repaired before ligating SMRTbell adapters (Pacific Biosciences). The resulting library was treated with an exonuclease cocktail to remove un-ligated DNA fragments, followed by two additional purification steps with AMPure XP beads (Beckman Coulter) and Blue Pippin (Sage Science) size selection to deplete SMRTbells < 7,000 bp. Libraries were then sequenced using P5 enzyme chemistry on the Pacific Biosciences RS-II platform to >200x genome-wide coverage.

### Complete genome assembly and finishing

PacBio SMRT sequencing data were assembled using a custom genome assembly and finishing pipeline (Altman et al. 2018). Briefly, sequencing data was first assembled with HGAP3 version 2·2·0 (Chin et al. 2013). Contigs with less than 10x coverage and small contigs that were completely encompassed in larger contigs were removed. Remaining contigs were circularized and reoriented to the origin of replication (*ori*) using Circlator (Hunt et al. 2015), and aligned to the non-redundant nucleotide collection using BLAST+ (Camacho et al. 2009) to identify plasmid sequences. In cases where chromosomes or plasmids did not assemble into complete circularized contigs, manual curation was performed using Contiguity (Sullivan et al. 2015). Genes were annotated using PROKKA (Seemann 2014) and visualized using ChromoZoom (Pak and Roth 2013) and the Integrated Genome Browser (IGB) (Nicol et al. 2009). Interproscan (Jones et al. 2014) was used to annotate protein domains and GO categories for annotated genes.

### Resolution of large genomic inversions

To resolve inversion events catalyzed by two prophage elements (*Staphylococcus phage* Sa1 and *Staphylococcus aureus* phage Sa5 with large (>40 kbp) nearly identical regions present in some of the assembled genomes, we developed a phasing approach that took advantage of unique variants present in each element. Raw (i.e. uncorrected) PacBio reads were first mapped to one of the repeat copies using BWA-MEM (Li 2013). Variants were then called with Freebayes (Garrison and Marth 2012), and high-quality single nucleotide variants with two distinct alleles of approximately equal read coverage were identified. Analogous to procedures used in haplotype phasing, we then determined which variant alleles were co-located in the same repeat element: if at ¾ of the raw reads containing a particular allele also encompassed distinct allele(s) of neighboring variant(s), the alleles were considered linked. In all cases this resulted in two distinct paths through the repeated prophage elements that were each linked to unique sequence flanking each repeat. We then used this information to correct assembly errors and identify *bona fide* inversion events between isolate genomes. Final verification of corrected assembly was performed by examining the phasing of the raw reads with HaploFlow (Bachmann et al. 2015).

### Phylogenetic reconstruction and molecular typing

Phylogenetic analyses were based on whole-genome alignments with parsnp (Treangen et al. 2014), using the filter for recombination. The VCF file of all variants identified by parsnp was then used to determine pairwise SNV distances between the core genomes of all strains. For visualization of the whole-genome alignments, isolate genomes were aligned using sibelia (Minkin et al. 2013) and processed by ChromatiBlocks (http://github.com/mjsull/chromatiblocks).

The multi-locus sequence type was determined from whole genome sequences using the RESTful interface to the PubMLST *S. aureus* database (Jolley et al. 2017). Typing of spa was performed using a custom script (https://github.com/mjsull/spa_typing). SCC*mec* typing was done using SCC*mec*Finder (Kaya et al. 2018). Changes to ACME and SaPI5 were determined using BLASTN and Easyfig. Presence or absence of genes in each locus was determined using BLASTX (Altschul et al. 1990) and a gene was considered to be present if 90% of the reference sequence was aligned with at least 90% identity. Prophage regions were detected using PHASTER. Each region was then aligned to a manually curated database of *S. aureus* phage intergrases using BLASTx to identify their integrase group.

### Annotation of antibiotic resistance determinants

Antibiotic resistance gene and variants were annotated by comparing to a manually curated database of 39 known *S. aureus* resistance determinants for 17 antibiotics compiled from literature. BLAST (Altschul et al. 1990) was used to identify the presence of genes in each isolate genome, with sequence identity cutoff ≥90% and an e-value cut-off ≤ 1e-10. Resistance variants were identified by BLAST alignment to the reference sequence of the antibiotic resistance determinant. Only exact matches to variants identified in literature were considered.

### Identification of transmissions

To establish similarity thresholds for complete genomes obtained from long read SMRT sequencing data we first examined baseline single nucleotide variant (SNV) distances between within each lineage. Median pairwise genome differences ranged from 101 SNVs for USA800 to 284 SNVs for USA100 (**Supplemental Fig. 8A**). We also examined the extent of divergence among 30 bacteremia isolate pairs collected within a span of one month to 1·4 years from individual patients. Pairwise distances for within-patient isolates were substantially lower than the median for each lineage (**Supplemental Fig. 8B-E**), consistent with persistent carriage of the same clone (Young et al. 2012; Von Eiff et al. 2001), with no more than 10 SNVs separating isolate pairs. Small (<5 bp) indels were more common than SNVs and mostly associated with homopolymer regions that can be problematic to resolve with third-generation sequencing technologies, indicating that they likely reflected sequencing errors. Notably, several patients showed variation between isolates collected within a span of several days (**Supplemental Fig. 8B-E);** indicative of intra-host genetic diversity. As such, we considered intra-host diversity and genetic drift in aggregate and set a conservative distance of ≤7 SNVs to define transmission events in our genome phylogeny.

### Identification of NICU outbreak subgroups

Changes between each outbreak isolate and the p133 reference isolate were identified using GWviz (https://github.com/mjsull/GWviz), which uses nucdiff (Khelik et al. 2017) to identify all genomic variants between pairs of strains. Nucdiff in turn uses nucmer to find alignments between two genomes and then identifies large structural rearrangements by looking at the organisation of nucmer alignments and smaller changes such as SNVs or indels by finding differences between the aligned regions. Briefly, raw PacBio reads were aligned back to each outbreak genome assembly using BWA-MEM (Li 2013). Provarvis was then used to detect and associate variants with PROKKA gene annotations, and to determine the number and proportion of raw reads supporting variants in each strain. Variants were selected for further delineation of outbreak subgroups if they were present in two or more isolate genomes and supported by at least ten raw reads in each genome, of which at least 75% confirmed the variant.

A graph of SNV distances between isolates was obtained from a multiple alignment of all outbreak isolates. The minimum spanning tree was then constructed using the minimum spanning tree functionality in the Python library networkX (https://networkx.github.io/).

### Identification of genetic variants unique to the NICU outbreak clone

To determine SNVs unique to the outbreak isolate the marginal ancestral states of the ST105 isolates were determined using RAxML (Stamatakis 2014) from a multiple alignment of all ST105s generated using Parsnp. We identified all SNVs that had accumulated from the most recent common ancestor of the outbreak strain and the closest related non-outbreak ST105, and the MRCA of all outbreak strains. SNVs causing nonsynonymous mutations or changes to the promoter region of a gene (defined as <500bp upstream of the start site) were plotted. Orthology was assigned using BLASTkoala (Kanehisa et al. 2016).

Core and accessory gene content in ST105 outbreak and non-outbreak strains was determined using ROARY. Genes found in more than two outbreak strains and less than 33% of the other ST105 genomes were then plotted along with select methylation data. Phylogenetic reconstruction of ST105 was performed using parsnp and the resulting tree and gene presence information was visualised using m.viridis.py (https://github.com/mjsull/m.viridis) which uses the python ETE toolkit (Huerta-Cepas et al. 2016).

### DNA methylation profiling

SMRT raw reads were mapped to the assembled genomes and processed using smrtanalysis v5·0 (https://www.pacb.com/products-and-services/analytical-software/smrt-analysis/).

Interpulse durations (IPDs) were measured and processed as previously described (Flusberg et al. 2010; Fang et al. 2012) to detect modified N6-methyladenine (^6m^A) nucleotides.

### RNA preparation and sequencing

For RNA extraction, overnight cultures in tryptic soy broth (TSB) were diluted (OD600 of 0·05), grown to late-log (OD600 of ~0·80) in TSB, and stabilized in RNALater (Thermo Fisher). Total RNA was isolated and purified using the RNeasy Mini Kit (Qiagen) according to the manufacturer’s instructions, except that two cycles of two-minute bead beating with 1 ml of 0·1 mm silica beads in a mini bead-beater (BioSpec) were used to disrupt cell walls. Isolated RNA was treated with 1 μL (1 unit) of Baseline Zero DNase (Epicentre) at 37°C for 30 min, followed by ribosomal RNA depletion using the Epicenter Ribo-Zero Magnetic Gold Kit (Illumina), according to the manufacturer’s instructions.

RNA quality and quantity was assessed using the Agilent Bioanalyzer and Qubit RNA Broad Range Assay kit (Thermo Fisher), respectively. Barcoded directional RNA-Sequencing libraries were prepared using the TruSeq Stranded Total RNA Sample Preparation kit (Illumina). Libraries were pooled and sequenced on the Illumina HiSeq platform in a 100 bp single-end read run format with six samples per lane.

### Differential gene expression analysis

Raw reads were first trimmed by removing Illumina adapter sequences from 3’ ends using cutadapt (Martin 2011) with a minimum match of 32 base pairs and allowing for 15% error rate. Trimmed reads were mapped to the reference genome using Bowtie2 (Langmead and Salzberg 2012), and htseq-count (Anders et al. 2015) was used to produce strand-specific transcript count summaries. Read counts were then combined into a numeric matrix and used as input for differential gene expression analysis with the Bioconductor EdgeR package (Robinson et al. 2010). Normalization factors were computed on the data matrix using the weighted trimmed mean of M-values (TMM) method (Robinson and Oshlack 2010). Data were fitted to a design matrix containing all sample groups and pairwise comparisons were performed between the groups of interest. P-values were corrected for multiple testing using the Benjamin-Hochberg (BH) method and used to select genes with significant expression differences (q < 0·05).

## Data Access

All genome data and assemblies are available on Genbank. Accession numbers are provided in **Table S1**.

## Acknowledgements

This research was supported in part by R01 AI119145 (H.v.B.), the Icahn Institute for Genomics and Data Science (A.K., E.S.), the NIAID-supported NRSA Institutional Research Training Grant for Global Health Research (T32 AI07647), the CTSA/NCATS KL2 Program (KL2TR001435, Icahn School of Medicine at Mount Sinai), and the New York State Department of Health Empire Clinical Research Investigator Program (Aberg, Icahn School of Medicine at Mount Sinai) (D.R.A.) and F30 AI122673 (T.R.P.). The funders had no role in study design, data collection and interpretation, or the decision to submit the work for publication. This research was also supported in part through the computational resources and staff expertise provided by the Department of Scientific Computing at the Icahn School of Medicine at Mount Sinai.

## Author Contributions

Methodology: M.J.S, D.A., H.v.B.; Data collection: M.J.S, D.A., K.C., B.C., E.W., T.R.P., G.D., M.L.S., Z.K., C.B., A.R., F.S., K.G., H.v.B.; Data curation: M.J.S, D.A., K.C., B.C.; Analysis: M.J.S, D.A., K.C.; Figures: M.J.S, D.A., K.C., H.v.B.; Writing – original draft: M.J.S, D.A., H.v.B.; Writing – review & editing: All authors; Study design: M.J.S, D.A., H.v.B., K.G.; Supervision: H.v.B., K.G.; Project administration: H.v.B., K.G.; Funding acquisition: H.v.B., D.A., T.R.P., A.K. E.S..

