## Supplemental Figures 1-8 for "Complete genome screening of clinical MRSA isolates identifies lineage diversity and provides full resolution of transmission and outbreak events"

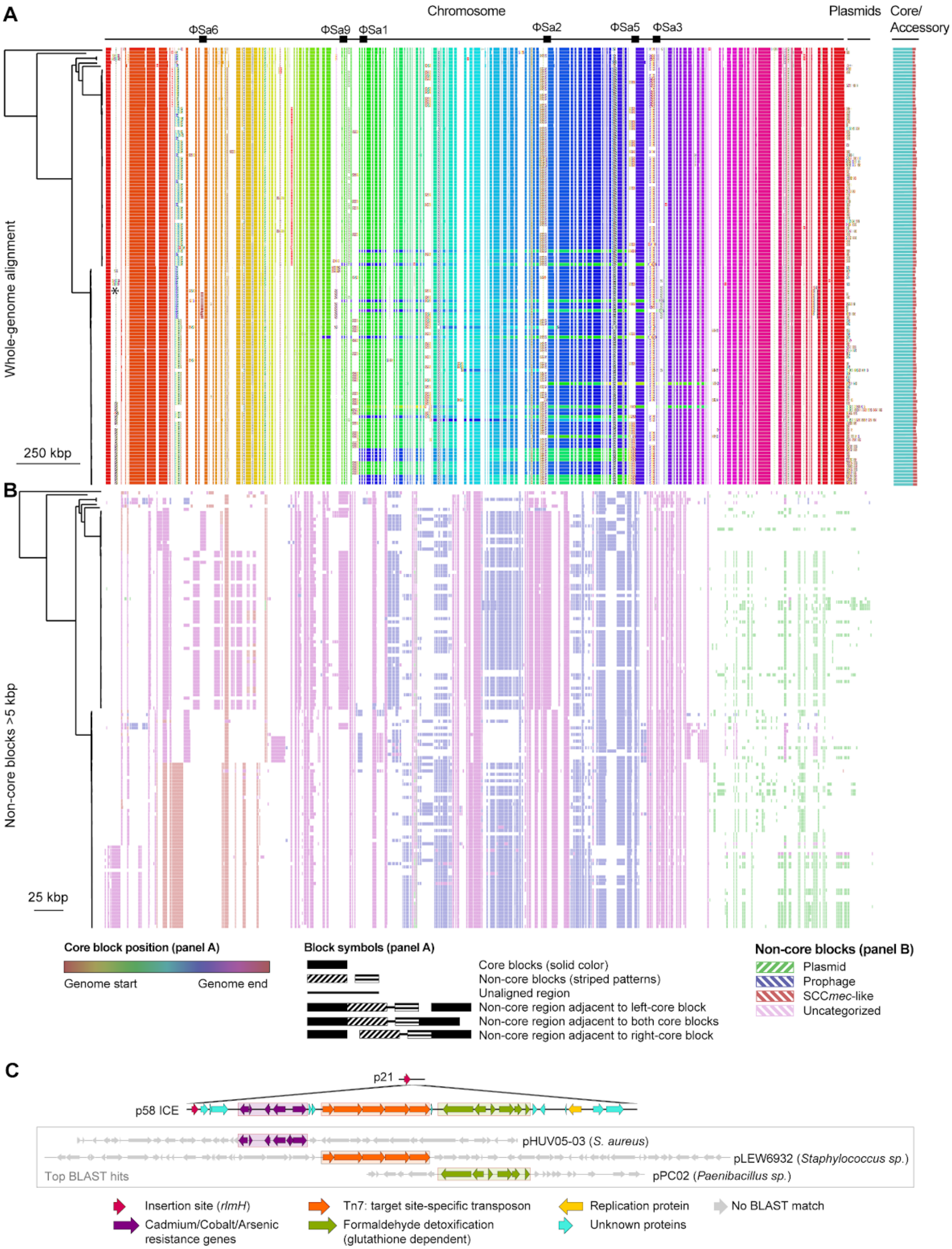
A
Chromosome
Plasmids
Core/ Accessory
ΦSa6
ΦSa9
ΦSa1
ΦSa2
ΦSa5
ΦSa3
Whole-genome alignment
250 kbp
B
Non-core blocks >5 kbp
25 kbp
Core block position (panel A)
Genome start
Genome end
Block symbols (panel A)
Core blocks (solid color)
Non-core blocks (striped patterns)
Unaligned region
Non-core region adjacent to left-core block
Non-core region adjacent to both core blocks
Non-core region adjacent to right-core block
Non-core blocks (panel B)
Plasmid
Prophage
SCCmec-like
Uncategorized
C
p21
p58 ICE
pHUV05-03 (S. aureus)
pLEW6932 (Staphylococcus sp.)
pPC02 (Paenibacillus sp.)
Top BLAST hits
Insertion site (rlmH)
Cadmium/Cobalt/Arsenic resistance genes
Tn7: target site-specific transposon
Formaldehyde detoxification (glutathione dependent)
Replication protein
Unknown proteins
No BLAST match

**Supplemental Figure 1. Multiple alignment of complete primary MRSA blood isolate genomes.** **A)** Maximum likelihood phylogenetic tree based on core genome SNVs of all primary MRSA blood isolates from 132 MSH inpatients (left), with a graphical representation of the complete genome alignment (right). Core syntenic blocks found in all strains are indicated by solid rectangles and accessory (non-core) syntenic blocks absent from at least one strain are indicated by patterned rectangles. Core blocks are colored according to the block location in each genome to highlight inversions (legend at bottom). Regions between the core blocks consist of accessory blocks (patterned rectangles) or unique sequence (solid black lines). These regions are placed equidistant between core blocks if the non-core region is found adjacent to the core block on either side. If the non-core region cannot be placed in between the core blocks to which it is adjacent (i.e. due to a structure of the genome being different to the reference) the region is attached arbitrarily to one of the core blocks to which it is adjacent. (see key below). Asterisk: location of the putative ICE element shown in C. **B)** Representation of all accessory genome blocks greater than 500 bp. Blocks are colored based on whether they originated from prophage, SCC*mec*, plasmid, or other uncategorized elements. A color key is included at the bottom. **C)** Example of a putative ICE element gained in isolate p21.

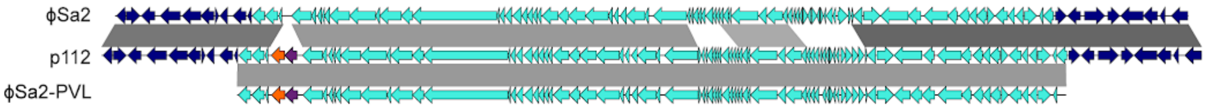


**Supplemental Figure 2. Acquisition of PVL in a USA100 isolate from p112.**
Diagram indicating a potential route for PVL acquisition in the p112 USA100 isolate (middle), by homologous recombination between a ΦSa2 prophage (top) and a ΦSa2-PVL prophage (bottom).

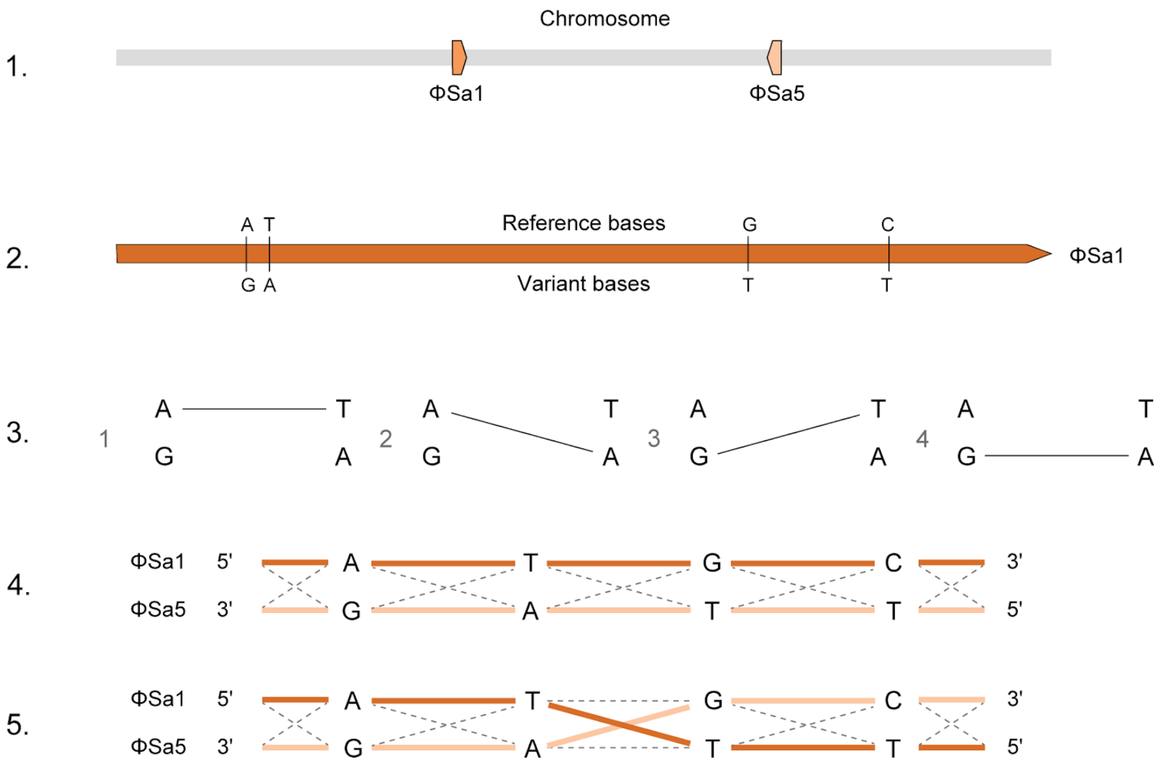


**Supplemental Figure 3. Resolution of inversions between ΦSa1 and ΦSa5 using PacBio raw reads.** The ΦSa1 and ΦSa5 prophages were first identified in the assembly using dot-plots or blast (#1). Next, BWA-MEM was used to align raw PacBio reads to one of the prophage sequences and SNVs that distinguish the two repeats were identified using freebayes (#2). Finally, the read coverage of each of the four possible combinations of linkage between the called SNVs were calculated, and a path from the unique regions at either ends was traced through the repeat (#3). If the identified path was consistent with the original assembly (i.e. #4) the sequence was left unchanged, otherwise the assembly was corrected (i.e. #5).

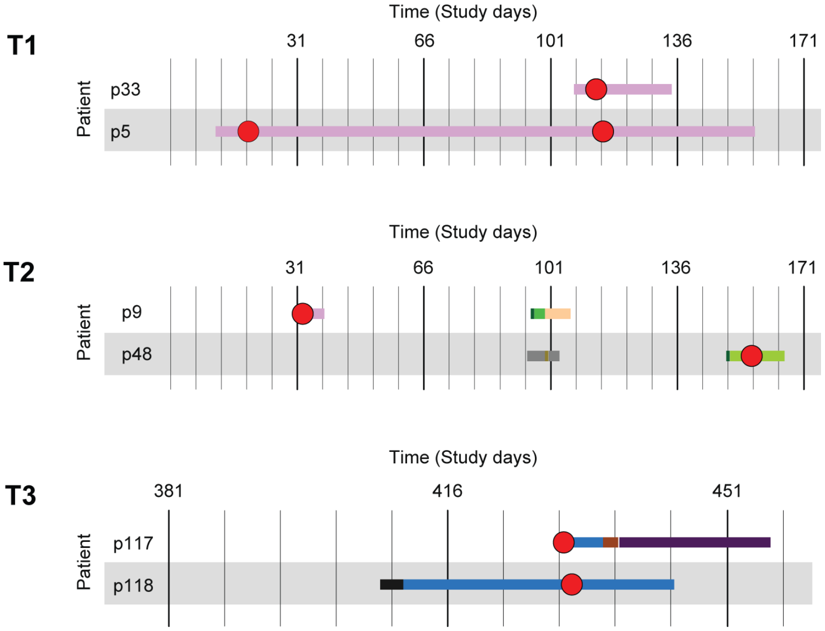


**Supplemental Figure 4. Transmission timelines for patients not involved in the NICU outbreak.** Overview of ward stays and isolates collected for the patients involved in transmission events T1, T2 and T3. Each row within a timeline corresponds to a patient, and admission periods are shown as horizontal bars where each ward is assigned a unique color. Red circles correspond to collection dates of positive blood cultures that were sequenced as part of the study.

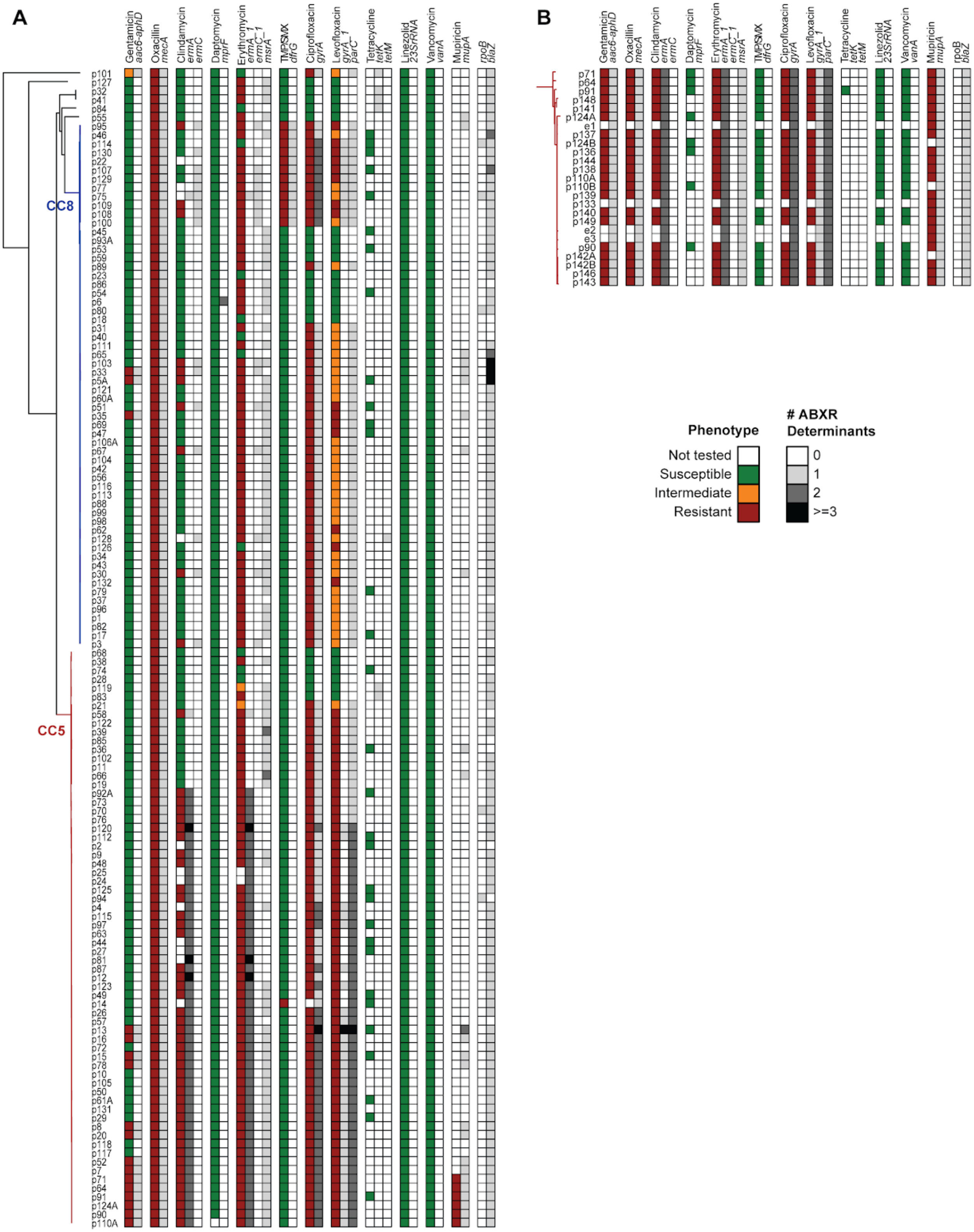


**Supplemental Figure 5. Overview of antibiotic resistance phenotypes and determinants.**
**A)** Results from antibiotic susceptibility testing (colored boxes) and genetic determinants (genes or mutations) identified in each genome (greyscale boxes) are shown for all primary MRSA blood isolates. Isolates are organized according to a maximum-likelihood phylogenetic tree based on SNV distances in core genome alignments. **B)** Same as A, but for the 25 isolates from adult and NICU outbreak cases.

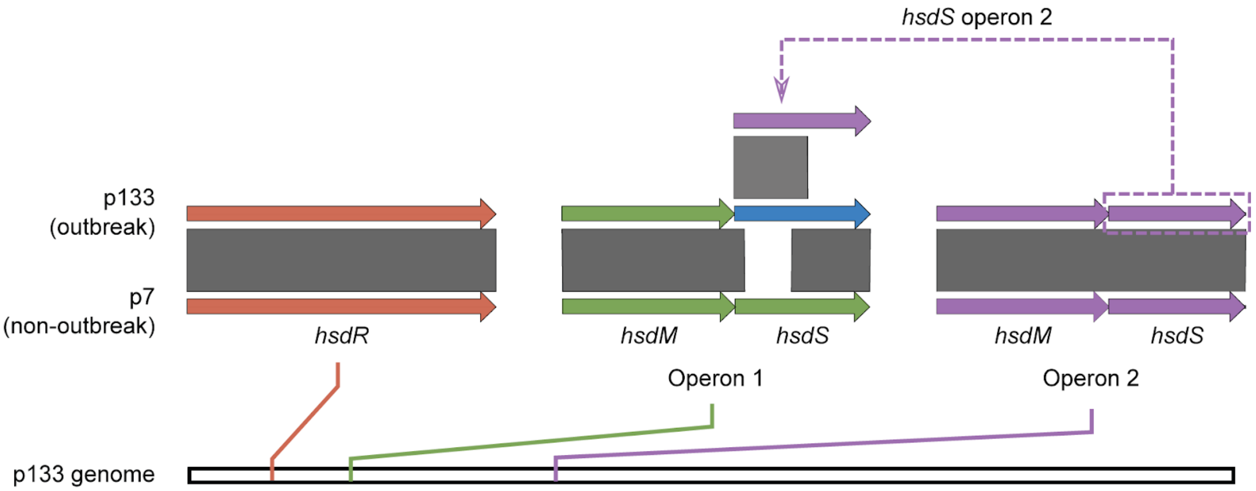


**Supplemental Figure 6. Recombination of the *hsdS* DNA binding domain in the NICU outbreak strain.** The type I restriction system in isolate p133 (representative of the outbreak strain) and p7 (representative of non-outbreak ST105 strains). The type I restriction system consists of a restriction protein (*hsdR*, red) and two operons (green and purple) with a modification (*hsdM)* and specificity gene (*hsdS*). Sequence similarity (determined by BLASTn) is shown by grey bars and the location of each loci in PS00004 is shown on the bottom. The *hsdS* gene in operon 1 of the outbreak strain (blue) differs from the typical ST105 isolate. The start of the gene is identical to the start of *hsdS* in operon 2 and the end of the gene identical to *hsdS* typical of operon 1.

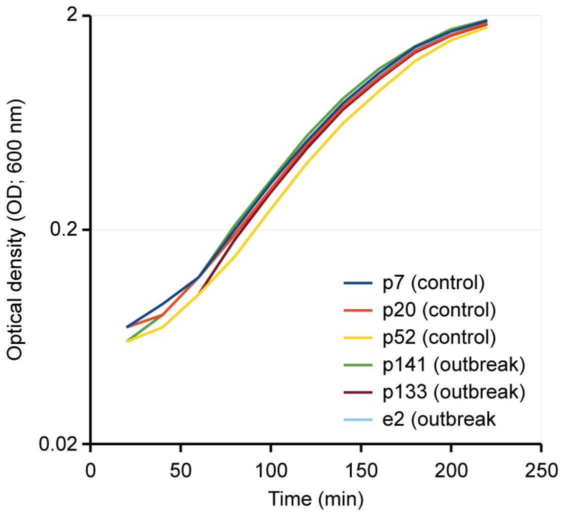


**Supplemental Figure 7. Growth curves of outbreak and control strains used for RNA-Seq.**
Representative growth curves of MRSA outbreak and closely related ST105 MSH control strains in TSB media.

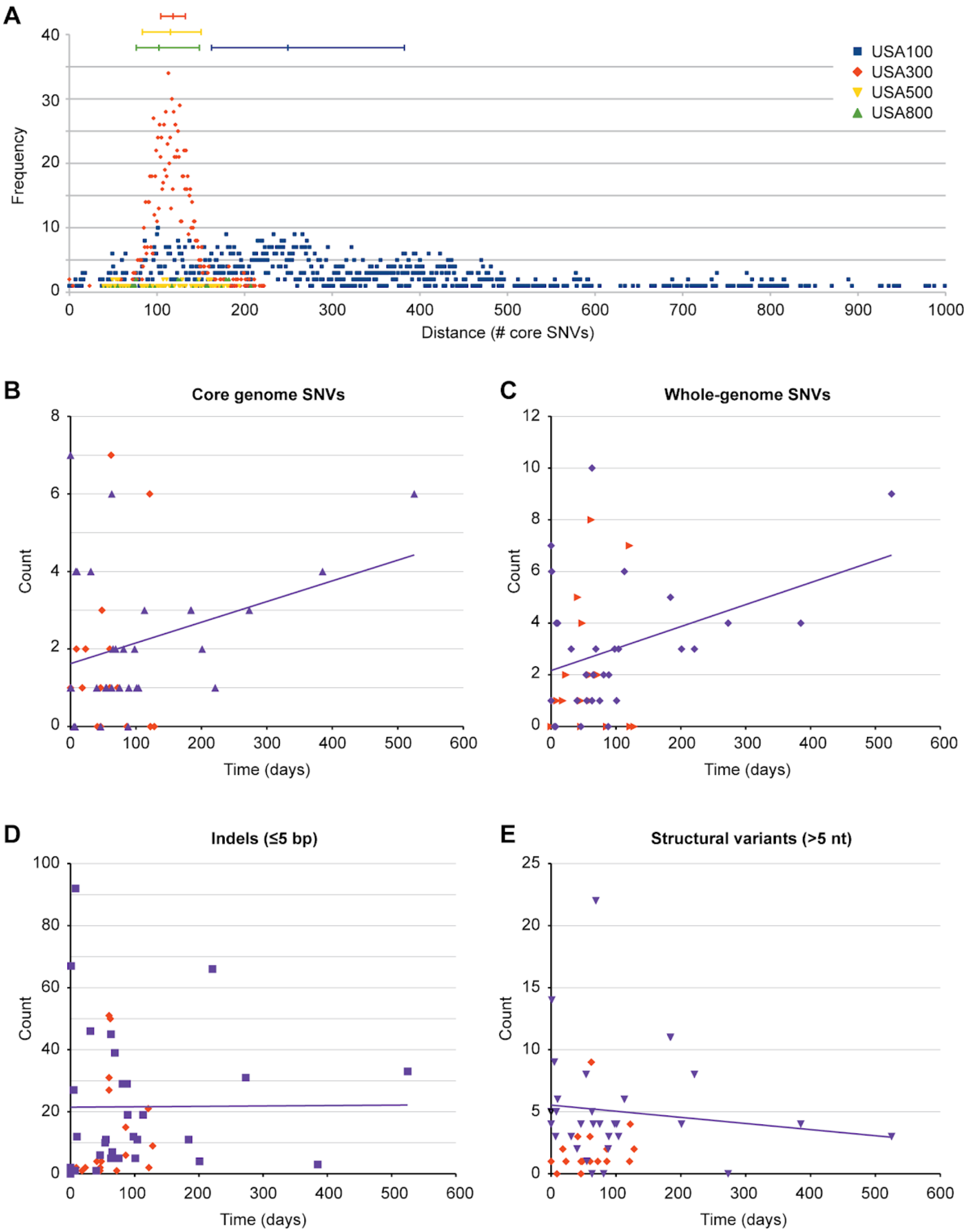


**Supplemental Figure 8. Genetic distances between isolates within MRSA lineages and between isolates obtained from single patients with persistent or recurrent infections.**

**A)** Frequency plot of SNV distances between all pairwise comparisons of genomes from primary MRSA bacteremia isolates within the USA100, USA300, USA500, and USA800 lineages. The median and interquartile range is indicated for each lineage above the plot. A bin size of 5bp was used. Below, the number of **B)** core-genome SNVs, **C)** whole-genome SNVs, **D)** indels ≤5 bp, and **C)** structural variants >5 bp between 30 isolate pairs obtained from the same patient, is plotted as a function of the time between collections. Data points from intra-patient isolate pairs are shown as purple triangles. Distances between NICU outbreak isolates are shown as orange diamonds, for comparison.
